# Harnessing *Escherichia coli* for bio-based production of formate under pressurized H_2_ and CO_2_ gases

**DOI:** 10.1101/2021.01.06.425572

**Authors:** Magali Roger, Tom C. Reed, Frank Sargent

## Abstract

*Escherichia coli* is gram-negative bacterium that is a workhorse of the biotechnology industry. The organism has a flexible metabolism and can perform a mixed-acid fermentation under anaerobic conditions. Under these conditions *E. coli* synthesises a formate hydrogenlyase isoenzyme (FHL-1) that can generate molecular hydrogen and carbon dioxide from formic acid. The reverse reaction is hydrogen-dependent carbon dioxide reduction (HDCR), which has exciting possibilities in bio-based carbon capture and storage if it can be harnessed. In this study, an *E. coli* host strain was optimised for the production of formate from H_2_ and CO_2_ during bacterial growth in a pressurised batch bioreactor. A host strain was engineered that constitutively produced the FHL-1 enzyme and incorporation of tungsten in to the enzyme, in place of molybdenum, helped poise the reaction in the HDCR direction. The engineered *E. coli* strain showed an ability to grow under fermentative conditions while simultaneously producing formate from gaseous H_2_ and CO_2_ supplied in the bioreactor. However, while a sustained pressure of 10 bar N_2_ had no adverse effect on cell growth, when the culture was placed at or above 4 bar pressure of a H_2_:CO_2_ mixture then a clear growth deficiency was observed. Taken together, this work demonstrates that growing cells can be harnessed to hydrogenate carbon dioxide and provides fresh evidence that the FHL-1 enzyme may be intimately linked with bacterial energy metabolism.

## 1. Introduction

In the current context of climate change, tackling carbon dioxide (CO_2_) emission levels by developing new technologies for carbon capture and utilisation is a global priority. Conversion of waste CO_2_ into marketable chemicals offers the possibility to achieve environmental sustainability and a circular economy while at the same time reducing atmospheric CO_2_ levels (Cheah et al., 2016; Schlager et al., 2017; Hepburn et al., 2019). Among the possible routes that can be considered for the capture of CO_2_, the direct hydrogenation of gaseous CO_2_ to aqueous formate (HCOO^−^) has gained attention since it offers a promising route to greenhouse gas sequestration, hydrogen (H_2_) transport and storage, and the sustainable generation of renewable chemical feedstock (Enthaler et al., 2010; Yuan et al., 2015; Yishai et al., 2016). Several chemical processes have been described that catalyse the reduction of CO_2_ to formate, but they require the use of expensive catalysts that generally operate under harsh conditions (Enthaler et al., 2010; Schaub and Paciello, 2011; Appel et al., 2013; Wang et al., 2015). In contrast, biological catalysts that are highly specific and work under milder conditions provide an attractive solution for the reduction of CO_2_ and the sustainable production of formate (Enthaler et al., 2010; Mellmann et al., 2016).

The gram-negative γ-Proteobacterium *Escherichia coli* is a facultative anaerobe that, under anaerobic conditions when using glucose as sole carbon and energy source, can perform a ‘mixed-acid fermentation’ that produces acetate, lactate, some succinate, and ethanol as end products. Mixed-acid fermentation also produces formate, which is often further disproportionated to H_2_ and CO_2_. The enzyme responsible for this is the formate hydrogenlyase (FHL-1) complex, which comprises a soluble catalytic domain containing a molybdenum- and selenocysteine (SeCys)-dependent formate dehydrogenase (FDH-H) module (encoded by the *fdhF* gene) linked by two [Fe-S]-cluster-containing proteins (encoded by *hycB* and *hycF*) to a nickel-dependent hydrogenase module (Hyd-3, encoded by *hycE* and *hycG*) (Sargent, 2016). The soluble catalytic domain of FHL-1 is anchored to the cytoplasmic membrane *via* two integral membrane subunits, encoded by *hycCD* genes, and under physiological fermentative conditions the FHL-1 forward reaction serves to detoxify formic acid accumulation and regulate environmental pH (Sargent, 2016).

FHL-1 can also operate in reverse as a H_2_-dependent CO_2_ reductase, HDCR (Pinske and Sargent, 2016). The yield of formate produced by the HDCR reverse reaction was initially low when carried out at ambient gas pressures, however the design of a laboratory-scale stirred tank reactor, which could be operated at precisely-controlled elevated gas pressures (up to 10 atmospheres), improved substrate solubility and gas transfer rates and led to a concomitant increase in the yield of the formate product (Roger et al., 2018).

The original HDCR experiments using pressurised gaseous substrates were conducted with pre-grown cell paste (i.e. non-growing cells) and carried out in the absence of any carbon or energy sources save for H_2_ and CO_2_ (Roger et al., 2018). In order to harness the HDCR activity for practical applications it would be desirable to enable *E. coli* to perform hydrogen-dependent CO_2_ reduction both under pressure and also during all active growth phases. In this study we used a multiscale bioengineering approach to tackle this issue by (i) optimising the host strain to produce FHL-1 under any growth regimen; (ii) attempting to remove bottlenecks in maturation and biosynthesis of FHL-1; and (iii) to chemically engineer a catalytic bias in favour of HDCR activity. To characterise the new bacterial strains and growth conditions, a laboratory-scale bioreactor dedicated for batch fermentation under pressurized H_2_ and CO_2_ was used. A strain that constitutively produced an engineered FHL-1 fusion protein was chosen that could perform the forward reaction and HDCR similar to the native enzyme. Incorporation of tungsten, in place of the native molybdenum, was shown to poise the engineered enzyme in the direction of hydrogen-dependent carbon dioxide reduction. Growth under a 10 bar nitrogen atmosphere had no effect on the growth rate of the engineered strain, however anaerobic growth rates were observed to decline under increasing H_2_/CO_2_ pressure as the cells were forced to perform the HDCR reaction. This work demonstrates that CO_2_ can be captured by FHL-1 activity in actively growing cells, and provides fresh insight in to the role of formate hydrogenlyase in *E. coli* energy metabolism.

## 2. Materials and Methods

### 2.1. Construction of bacterial strains

This work was based on *E. coli* K-12 MG059e1 which carries a *hycE*^HIS^ allele on the chromosome (McDowall et al., 2014). Strains constructed and employed in this study are listed in Table 1. Gene deletions were carried out by transduction with bacteriophage P1 (Thomason et al., 2007). Strains from single-gene knockout ‘Keio’ collection of the non-essential genes of the *E. coli* K-12 BW25113 were used as donors. Deleted genes were *hyaA*: catalytic subunit of Hyd-1, *hybC*: catalytic subunit of Hyd-2, *pflA*: pyruvate formate lyase activating protein; *fdhE*: Tat-dependent formate dehydrogenases accessory protein; *iscR*: transcriptional repressor of *isc* operon; *metJ*: transcriptional repressor of *met* regulon. Once target genes were replaced with a kanamycin-resistance gene,the resultant strains were transformed with plasmid pCP20 (Baba et al., 2006) before the plasmid carrying the resistance cassette was cured at 42°C. Gene deletions and antibiotic resistance loss were confirmed by PCR. The strain producing the FhlA^E183K^ variant was obtained by introducing a single base pair substitution in the *fhlA* gene. First, the 500 bp upstream and downstream of *fhlA* gene was amplified by PCR and assembled in pMAK705 (Hamilton et al., 1989). The mutant allele was then incorporated by QuickChange PCR and then transferred on to the chromosome by homologous recombination (Hamilton et al., 1989). Strain producing FdhF as an N-terminal fusion protein to HycB joined by a linker sequence containing a hemagglutinin (HA) tag flanked by three glutamines on each side was constructed as previously reported (McDowall et al., 2015). To upregulate expression of ϕ *fdhF*^HA^-*hycBCDEFGHI*, the synthetic T5 promoter region from pQE60 (Qiagen), the *E. coli proD* promoter, and the promoter of *tatA* and *ynfE* genes were amplified and cloned into pMAK705 as an EcoRI-BamHI fragment. Finally, the 500 bp upstream of *hycA* and downstream of the ϕ *fdhF*^HA^::*hycB* were cloned in the same vectors as a KpnI-EcoRI and BamHI-HindIII fragments, respectively.

**Table 1:**
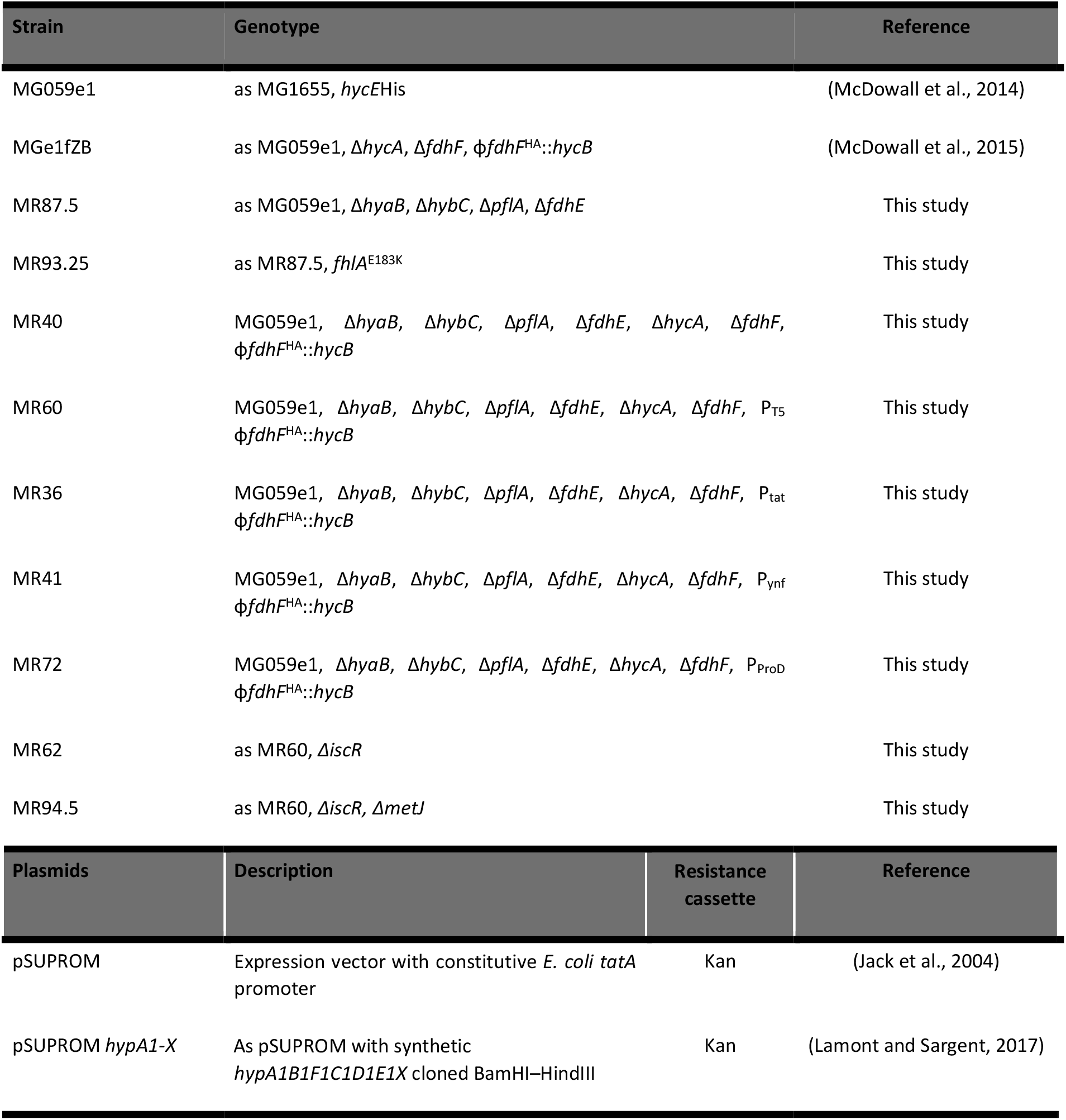
Strains and plasmids used in this study.

### 2.2. Protein purification and analytical methods

Cells that were grown under anaerobic fermentative conditions (5 L) were harvested by centrifugation and suspended with lysis buffer containing 50 mM Tris HCl pH 8.0 with 10 μg/mL DNase I (Sigma), 50 μg/mL lysozyme (Sigma), and a protease inhibitor cocktail (<u>Roche</u>) at 1 g wet cell weight per 10 mL of buffer. Cells were lysed using a high-pressure cell homogeniser (Homogenising System Ltd) at 1,000 bar before unbroken cells and debris were removed by centrifugation. Membrane proteins were solubilised for 1.5 hours at room temperature by adding N-Dodecyl-β-D-Maltopyranoside (DDM) 1 % (w/v) directly to the crude extract. Then, the solubilised fraction was loaded onto a 5 mL HisTrap HP column (GE Healthcare) that had been equilibrated in 50 mM Tris∙HCl (pH 7.5), 150 mM NaCl, 50 mM imidazole, 0.02% (w/v) DDM. Bound proteins were eluted with a 6-column volume linear gradient of the same buffer containing 500 mM imidazole. Fractions (10 μL) were analysed by SDS-PAGE using the method of Laemmli (Laemmli, 1970) and stained with Instant Blue. Fractions containing FHL-1 components were pooled and concentrated in a Vivaspin (Millipore Inc.) filtration device (50 kDa molecular weight cut off). Protein concentration in each fraction was determined using DC protein determination kit for detergent-containing samples. For Western immunoblotting, samples were first separated by SDS-PAGE before transfer to nitrocellulose (Dunn, 1986). Nitrocellulose membranes were developed using a mouse anti-HA monoclonal antibody (Invitrogen), and a goat anti-Mouse HRP-conjugated secondary antibody (Bio-Rad). Nitrocellulose membranes were analysed with the ImageQuant Las 4000 imager (GE Healthcare).

### 2.3. Whole cell catalysis of H2/formate production at ambient pressure

Cells were grown under anaerobic fermentative condition in rich LB medium containing 0.4 % (w/v) glucose. When mentioned, the medium was supplemented with 0.2 % (w/v) and/or 1 μmol.L^−1^ sodium tungstate. The culture was harvested by centrifugation (Beckman J6-MI centrifuge) for 30 min at 5000 g and 4°C. The cell paste was washed twice in 20 mmol.L^−1^ 3-(N-morpholino)propanesulfonic acid (MOPS) buffer, pH 7.4, before the cell pellet was suspended in the same buffer and volume was adjusted to reached OD at 600nm of 0.5 (~0.125 g_CDW_). For the FHL/Fwd activity, 0.2 % (w/v) sodium formate was added to the cell suspension and 5 mL of washed whole cells were transferred into Hungate tubes that were flushed with nitrogen for 5 min. For the FHL-1 reverse reaction, 3 mL of washed-whole cells were transferred into Hungate tubes. Tubes were flushed with N_2_ for 5 min, then with H_2_ for another 5 min before 5 mL CO_2_ were added to the tubes. The cells were incubated at 37°C for 20 hours. H_2_ content in the headspace and formate content in the cell suspension were determined by gas chromatography (GC) and High-Performance Liquid Chromatography (HPLC).

### 2.4. Pressurised bioreactor culture conditions

The cultures were performed in a commercially-available bioXplorer^®^ 400P system (HEL Ltd, UK). The working liquid volume was 250 mL. The bioreactor was set with three gas inlets H_2_, CO_2_ and N_2_ (BOC, UK). Each gas inlet was controlled by a gas mass flow controller. The bioreactor was equipped with pH, dissolved oxygen (DO), temperature and pressure controllers. The WinIso^®^ software handled the on-line monitoring and control systems of the reactor. Temperature was maintained at 37 °C. The pH was maintained at 7.0 by addition of a 5 mol.L^−1^ NaOH solution. The bioreactor containing LB rich medium was first autoclaved before 0.8 % (w/v) glucose and 1 μmol.L^−1^ sodium tungsten were added prior to inoculation. A 10% (v/v) culture grown in LB rich medium under aerobic conditions for at least 16 hours were used to inoculate the bioreactor. After inoculation, O_2_ present in the medium is rapidly consumed by the culture as observed via the in-line O_2_ electrode. After this point the H_2_, CO_2_ and/or N_2_ gases were sparged through the medium at a gas flow rate of 50 mL.min^−1^.

### 2.5. Analytical methods

Bacterial growth was monitored by following the OD at 600nm and the biomass yield was estimated from the OD at 600nm of the culture and the assumption that 1 L culture with an OD600 of 1 contains 0.25 g_CDW_. Metabolite analysis and quantification were determined by HPLC using a an UltiMate 3000 uHPLC system (Thermo) equipped with an Aminex HPX-87H column (BioRad) using a RefractoMax521 refractive index detector (Thermo) and a Variable Wavelength Detector-3100 at 210nm (Thermo). Typically, samples of 10 μL that were previously clarified through 0.2 μm filters were applied to the column equilibrated in 5 mmol.L^−1^ H_2_SO_4_ with a flow of 0.8 mL.min^−1^ at 50°C. Hydrogen in the headspace was quantified using GC-2014 gas chromatograph (Shimadzu) equipped with a capillary column and thermal conductivity detector. Typically, 1 mL samples were collected using a syringe with Luer lock valve (SGE) and used to manually fill a 500 μL loop. Nitrogen was used as carrier gas with 25 mL.min^−1^ of gas flow rate.

## 3. Results and Discussion

### 3.1. Harnessing FHL-1 expression by genetic engineering

The first obstacle to overcome for exploiting *E. coli* FHL-1 as a carbon fixing technology is the natural expression regime of the enzyme, which is geared towards environmental conditions favouring the forward reaction. Thus, normal FHL-1 biosynthesis is controlled by the presence of formate, acidic pH and the σ^54^ transcription factor (Birkmann et al., 1987; Schlensog et al., 1989; Rossmann et al., 1991; Leonhartsberger et al., 2002). The expression of the *fdhF* gene and *hyc* operon is also coordinated and regulated by a formate-responsive transcriptional regulator FhlA (Schlensog and Böck, 1990; Schlensog et al., 1994) and the repressor HycA (Böhm et al., 1990; Sauter et al., 1992). In the presence of formate, FhlA is activated and induces the transcription of the FHL-1 complex *via* σ^54^. Hence, numerous studies have demonstrated that the overexpression/mutation of the FhlA transcriptional activator, along with the deletion of *hycA* gene, often resulted in improving bio-H_2_ production due to up-regulation of FHL-1 (Penfold et al.; Korsa et al., 1997; Yoshida et al., 2005; Sanchez-Torres et al., 2009). Moreover, it was also shown that these mutations combined with other mutations aiming at inactivating competing pathways can have synergistic effects (Bisaillon et al., 2006; Yoshida et al., 2006; Maeda et al., 2008; Mathews et al., 2010). Thus it is clear that strategies to remove formate control of FHL-1 biosynthesis are needed in order to produce active FHL-1 under all growth regimens.

First, an *E. coli* strain (MR87.5) was constructed in which other hydrogenases (Δ*hyaB*, Δ*hybC*) and potential formate utilization pathways (Δ*pflA*, Δ*fdhE*) were inactivated (Table 1). As previously observed in the context of bio-H_2_ production (Adams, 1990; Frey, 2002), mutant strains unable to synthetize active pyruvate formate lyase (Δ*pflA*), which normally generates formate from pyruvate during fermentation, should only produce FHL-1 when formate is supplemented to the growth medium. This phenotype was observed here for strain MR87.5 (*hycE*^His^, Δ*hyaB*, Δ*hybC*, Δ*pflA*, Δ*fdhE*) (Figure 1B). Here, the H_2_-dependent CO_2_-reductase (HDCR, reverse FHL-1) activity was only observed in MR87.5 after the cells have been pre-grown in exogenous formate (Figure 1B).

**Figure 1:**
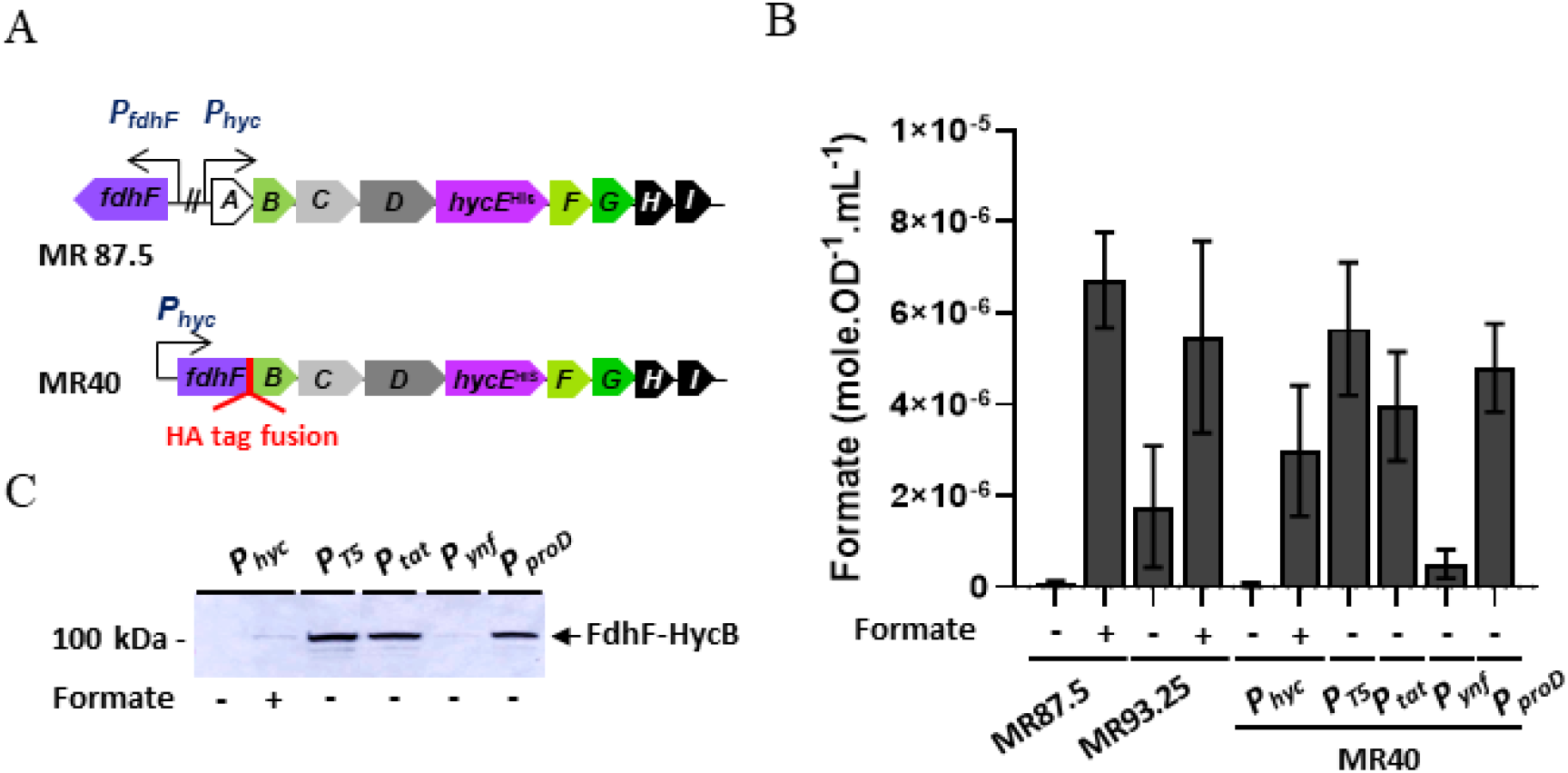
Harnessing FHL-1 production and HDCR activity by genetic engineering. **(A)** Genetics of the *E. coli* FHL system in a parental (MR87.5) and engineered (MR 40) strains context. The *hyc* operon is encoded at 61 min on the genome, while *fdhF* is located at 91 min. Both transcriptional units are regulated by the formate-responsive transcriptional activator *fhlA*. For the construction of the engineered *hyc* operon, the *fdhF* gene was deleted in this genetic background, and the *hycA* gene coding for a transcriptional repressor was replaced by a new version of the *fdhf* gene fused with *hycB via* a HA epitope tag according to (McDowall et al., 2015). Alternatively, the native promoter P*hyc* was replaced by synthetic promoters P_T5_, P_*tat*_, P_*ynf*_ and P_proD_. **(B)** End-point hydrogen-dependent carbon dioxide reduction (formate production) assays using cells pre-grown under anaerobic fermentative conditions ^+^/-0.2 % (w/v) formate. **(C)** The production of the FdhF-HycB fusion protein was assayed by Western immunoblot using an antibody against the HA epitope in the fusion linker sequence.

Next, the FhlA regulator was specifically targeting for mutagenesis. Among the FhlA variants characterised so far, it was shown that FhlA with amino acid replacements E183K not only resulted in improved *hyc* transcription, but it also conferred a constitutive phenotype apparently blind to formate concentrations (Korsa et al., 1997). In this work, the MR87.5 strain was further modified by the inclusion of an FhlA^E183K^ allele to give *E. coli* strain MR93.25 (Table 1). This new strain demonstrated some HDCR activity when initially cultured in the absence of exogenous formate (Figure 1B), however, surprisingly, HDCR activity in the FhlA E183K variant remains strongly inducible by pre-growth in external formate (Figure 1B).

Harnessing production of an intact FHL-1 enzyme is further complicated by the fact that the formate dehydrogenase and hydrogenase genes are located at separate loci on the chromosome. In an initial attempt to improve co-production of the entire FHL complex, a previous study engineered an *E. coli* strain in which the FDH-H moiety was physically tethered to HycB resulting in the production of a fully assembled and functional complex (McDowall et al., 2015). Keeping with this strategy here, the *fdhF* gene was first deleted from the parental strains before the *hycA* gene encoding a transcriptional repressor was replaced by a version of the *fdhF* gene fused to *hycB* using a HA epitope tag (Figure 1A). This new strain (MR40) was then further modified by the inclusion of alternative transcriptional promoter regions upstream of the ϕ*fdhF*::*hycB* fusion allele (Table 1). The promoters T5, *proD*, *tatA* and *ynfE* were chosen as various examples of strong, constitutive or anaerobically-induced promoter sequences. The four new strains carrying these promoters (Table 1) were then analysed for the production of the FdhF^HA^-HycB fusion protein by Western immunoblotting against the HA tag (Figure 1C) and for HDCR activity using intact whole cells (Figure 1B). As shown in Figure 1, when the expression of FHL-1 was left under the control of what remains of the native *hyc* promoter (P*hyc*) in MR40, the cells exhibited no HDCR activity when cultivated in the absence of formate and no protein could be detected by Western Blot using an antibody raised against the HA epitope tag. Moreover, this strain yielded only low levels of protein when cells were grown with extra formate in the medium (Figure 1C). As a result, the HDCR activity was only partially restored when formate was supplemented in the growth medium (Figure 1B). Next, the synthetic promoter constructs were tested. Among the promoters screened, T5 is a strong promoter that allows constitutive expression in the absence of the LacI repressor and the promoter region upstream of *ynfE* gene which was proposed to be regulated by FNR and shown to be highly upregulated under anaerobic conditions (Kang et al., 2005). A promoter, termed *proD*, which is a constitutive that exhibits high transcription rates independently of the genetic context (Davis et al., 2011) was also tested. The MR60 strain, which was engineered with a strong T5 promoter upstream of the ϕ*fdhF*^HA^*::hycB* allele fusion, showed the most the convincing protein production yield in the absence of exogenous formate (Figure 1C) and the highest HDCR activity (Figure 1B). Interestingly, formate production yield in mutant strains never exceeded that observed using the parental strain (Figure 1B) suggesting that production of the enzyme complex ultimately is not currently the limiting factor in formate production.

### 3.2. Exploring optimisation of FHL-1 cofactor biosynthesis and insertion

Maturation of FHL-1 is a multi-step process that depends on accessory proteins involved in the biosynthesis of the [Fe-S] clusters, the Molybdenum cofactor (also termed as MoCo) of the formate dehydrogenase, and the [NiFe] active site cluster (*i.e.* Ni–Fe–CO–2CN^−^) of the hydrogenase. Therefore, it can be hypothesized that the amount of functional enzyme under any overproduction regime is limited by cofactors biosynthesis machineries. Previous strategies to stimulate hydrogenase expression and activity has involved the deletion of the *iscR* gene in *E. coli* (Akhtar and Jones, 2008; Jaroschinsky et al., 2017). In *E. coli*, *iscR* encodes a transcriptional master regulator that operates as a feedback loop to repress transcription of the *iscRSUA* operon, as well as more than 40 other genes with known or unknown physiological roles (Schwartz et al., 2001; Giel et al., 2006, 2013), which forms the ISC system involved in providing [Fe-S] clusters to the hydrogenase (Pinske and Sawers, 2012; Pinske et al., 2013). Here, a version of the MR60 strain carrying a Δ*iscR* allele (*E. coli* strain MR62) was constructed (Table 1).

Attention was next given to the MoCo biosynthesis pathway. The biosynthesis of MoCo in prokaryotes is a highly conserved and complex mechanism involving a series of accessory proteins and co-substrates(Magalon et al., 2011; Iobbi-Nivol and Leimkühler, 2013; Leimkühler et al., 2016). Here, we focused on the synthesis of the S-adenosyl methionine (SAM) radical which plays a critical role in the first step of the pathway(Sofia et al., 2001; Hänzelmann and Schindelin, 2006). The genes involved in SAM biosynthesis in *E. coli* are located on different transcriptional units on the chromosome and are referred to as the ‘*met*’ regulon(Weissbach and Brot, 1991). Interestingly, the *met* regulon is regulated by a master transcriptional regulator, encoded by the *metJ* gene that operates as a feedback loop to repress transcription(Weissbach and Brot, 1991; Nakamori et al., 1999). Moreover, the MoCo biosynthesis pathway is intimately connected to [Fe-S] clusters and the ISC machinery (Yokoyama and Leimkühler, 2015; Mintmier et al., 2020). Therefore, the MR62 strain carrying Δ*iscR* was further engineerd by the inclusion of a Δ*metJ* deletion to yield strain MR94.5 (Table 1).

Finally, it was considered worthwhile to attempt to boost the [NiFe] cofactor biosynthesis capability in the strains, and to do this cells were transformed with a multi copy vector carrying a synthetic version of the *hypA1B1C1D1E1X* operon from *Ralstonia eutropha* (*Cupriavidus necator*). This strategy has proven to be successful in recent studies of hydrogenase overproduction (Lamont and Sargent, 2017; Beaton et al., 2018).

Strains with engineered cofactor biosynthesis pathways were analysed for the FHL-1 forward reaction − H_2_ production. As shown Figure 2, H_2_ production initially decreased in the MR40 parent strain carrying the ϕ*fdhF*^HA^*::hycB* allele fusion, while the incorporation of a T5 promoter upstream of the fusion in the MR60 strain restore the activity to native levels. The subsequent deletions of the *iscR*, metJ or inclusion of extra [NiFe] cofactor accessory genes added no material improvements to FHL-1 activity. (Figure 2). This clearly shows that, under these growth conditions, the metal cofactor biosynthesis, insertion and maturation pathways of the enzyme were not a limiting factor.

**Figure 2:**
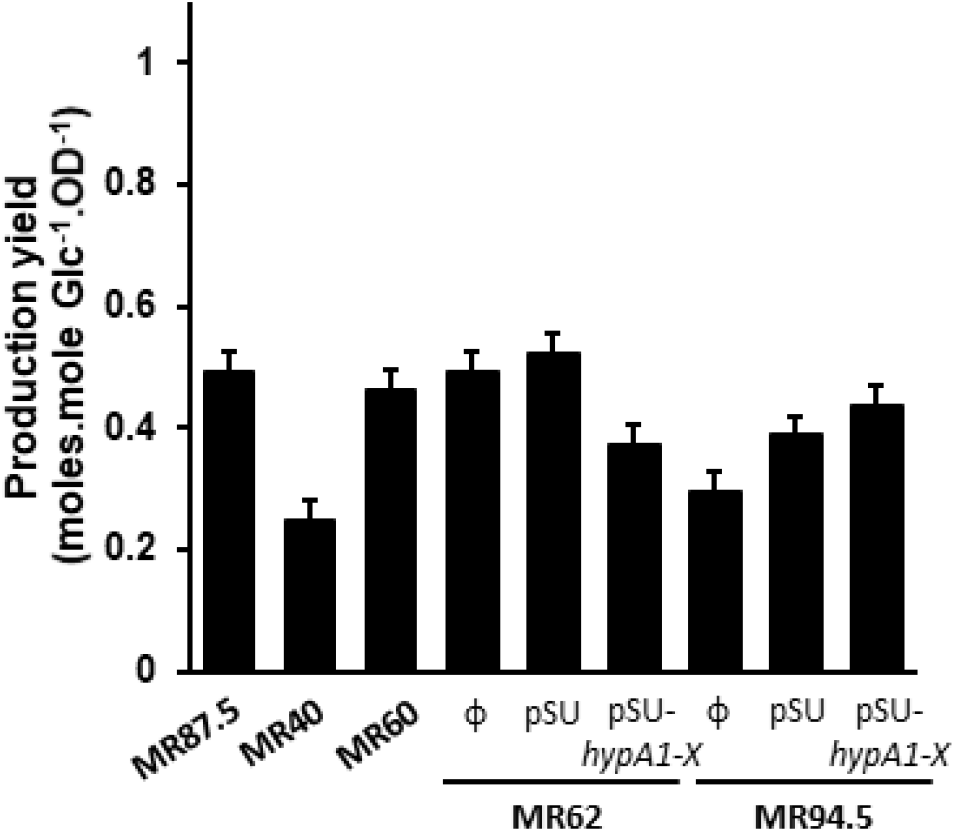
Optimising FHL-cofactor maturation by genetic engineering. *E. coli* strains (Table 1) were grown under anaerobic fermentative conditions in rich medium supplemented with 0.4 % (w/v) glucose and 0.2 % (w/v) formate for 20 hours incubation at 37°C. The FHL-1 activity was assayed by measuring H_2_ content in the gas phase by GC (n = 3).

### 3.3. Biasing HDCR activity by biochemical engineering of the formate dehydrogenase

One major obstacle to consider is the thermodynamics and reversibility of the FHL system. Clearly, it would be desirable to minimise any tendency towards the forward reaction while HDCR activity is being harnessed. Under ambient conditions (at 1 atmosphere and room temperature), the FHL-1 formate dehydrogenase was shown to reduce CO_2_ but at considerably lower rates than the forward reaction of formate oxidation (Bassegoda et al., 2014; McDowall et al., 2015; Pinske and Sargent, 2016). Indeed, the activities of metal dependent formate dehydrogenases for formate oxidation and CO_2_ reduction vary greatly depending on the metal co-factor composition (Molybdenum or Tungsten-W) and its coordination (either by Cys or SeCys amino acid side-chains) (Maia et al., 2015; Niks and Hille, 2019). Overall, W-containing formate dehydrogenases have been suggested to be more efficient at reducing CO_2_ than the Mo-containing variants because of the lower midpoint potential of the active site cofactor (Maia et al., 2017). Moreover, it was reported that a few enzymes can be active with either Mo or W (Buc et al., 1999; Stewart et al., 2000; Brondino et al., 2004). Thus, it was considered here whether FHL-1 could be produced as a variant containing W instead of Mo in the formate dehydrogenase active site.

First, MR60 *E. coli* cells were grown under anaerobic conditions using a medium supplemented with or without extra tungstate ions. The FHL-1 complex was then purified as previously described (McDowall et al., 2014, 2015) and the metal content of the purified enzyme determined by ICP-MS. As shown Table 2, the native purified enzyme isolated *via* a His-tag on Hyd-3 contains clearly detectable amounts of nickel ions in a ratio of 1:0.4 with molybdenum and 1:0.01 with tungsten. Thus the native enzyme contain essentially no tungsten relative to nickel. Interestingly, when the growth medium contains tungsten salts, the purified enzyme is seen to contain nickel in a ratio of 1:0.0006 with molybdenum and 1:0.7 with tungsten, suggesting that tungsten can be erroneously incorporated into the enzyme when supplied in excess.

**Table 2:** Ratio of purified FHL-1 active site metal content as determined by ICP-MS.

| Growth Conditions | <sup>98</sup>Mo :<sup>58</sup>Ni | <sup>182</sup>W :<sup>58</sup>Ni |
| --- | --- | --- |
| LB medium | 0.4049 | 0.0006 |
| LB + 1 mmol.L<sup>-1</sup><br>Na<sub>2</sub>WO<sub>4</sub> | 0.0126 | 0.7447 |

Next, the cells growing in the presence of increasing quantities of tungsten salts were analysed for both forward and reverse reactions of FHL-1 (Figure 3). Both FHL-1 forward and HDCR activities tended to decrease as the concentration of tungstate ions increased in the growth medium. However, the profiles of inhibition were strikingly different. Notably, at 1 μM tungstate in the growth medium, a 50 % loss of FHL-1 forward (H_2_-production) activity was observed, while the same cells retained full HDCR activity under these conditions. This result strongly suggests that the substitution of the Mo atom at the active site of FdhF by a W atom can either shift the catalytic bias towards CO_2_ reduction or simply inhibit the forward reaction. Nevertheless, for the first time here we show that turning the FdhF from *E. coli* into a W-variant could deliver a unidirectional HDCR that promotes CO_2_ reduction even under ambient pressures and high product concentrations.

**Figure 3:**
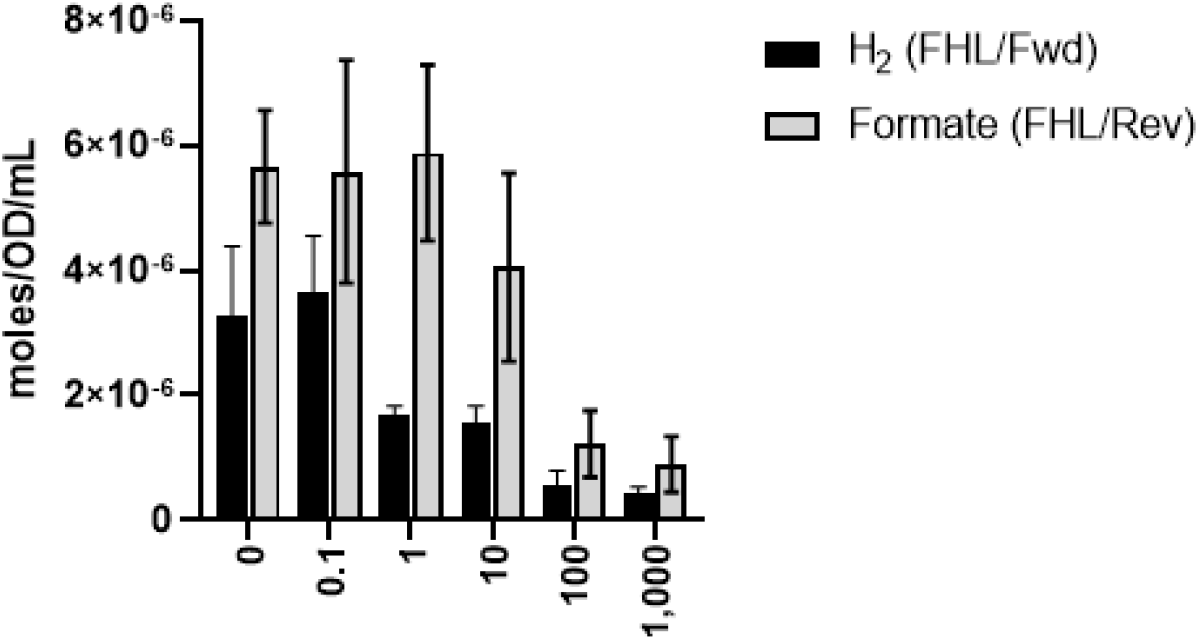
Chemical engineering of the formate dehydrogenase. *E. coli* MR60 (Table 1) was pre-grown under anaerobic fermentative conditions in rich medium containing 0.4 % (w/v) glucose supplemented with variable amounts of tungstate salts (0-1000 μM final concentration). Cells were harvested in stationary phase, washed twice with a buffer at pH 7.4 and incubated with 20 % (w/v) formate under nitrogen atmosphere or a 50/50 H_2_:CO_2_ atmosphere. Finally, FHL-1 forward activity and HDCR activity were assayed following 20 hours incubation at 37 °C by GC and HPLC, respectively.

Interestingly, both forward and reverse activities were lost when cells were grown with the highest concentration of tungstate ions (1 mmol/L). It has been shown that the expression of *fdhF* and *hyc* is regulated by molybdate concentration in the cell through the action of the transcriptional regulator, ModE(Rosentel et al., 1995), however the effect of W on *fdhF* and *hyc* operon expression, or cofactor biosynthesis, remains largely unexplored in this context. Previous studies showed that the incorporation of Mo or W at the active site of formate dehydrogenases in *Desulfovibrio* species is regulated not only by different selectivities in metal incorporation but also at the level of gene expression (Brondino et al., 2004; da Silva et al., 2011; Mota et al., 2011).

### 3.4. Developing a bioprocess for CO_2_ hydrogenation by E. coli throughout bacterial growth

#### 3.4.1 Formate production by the optimised strain of *E. coli* in batch fermentation at ambient pressure

The ultimate goal of this research is to generate a host stain, and define some growth conditions, that will perform HDCR throughout the growth phase. Next, we employed a bioXplorer P400 laboratory-scale bioreactor with a gas sparging system to allow a constant and efficient delivery of H_2_, CO_2_ and/or N to the culture. The engineered MR60 strain, which can only generate formate *via* engineered FHL-1, was chosen for initial trials and was grown in the presence of tungstate salts to maintain HDCR activity. Following inoculation, the oxygen present in the growth medium was observed to be rapidly consumed by the bacteria. When the oxygen dropped down to 0 %, only then were H_2_ and CO_2_ sparged through the cell culture at 50 mL.min^−1^. In this first experiment, no overpressure was applied. Initially, a concomitant with a drop of pH in the growth medium was observed as CO_2_ was added. Hence, sodium hydroxide was automatically pumped into the growth medium to maintain the pH at 7.0 (Supp. Fig. SI S1). Bacterial growth was observed in the first 8 hours of the run, until glucose was fully consumed, reaching a maximum turbidity OD_600nm_ of 1.8 (*i.e.* 0.45 g_CDW_.L^−1^) (Figure 4). Under these conditions, a maximum of ~ 8 mmol.L^−1^ formate was produced after 24 hours with a maximum rate of formate production of 2.4 mmole.L^−1^.h^−1^.

**Figure 4.**
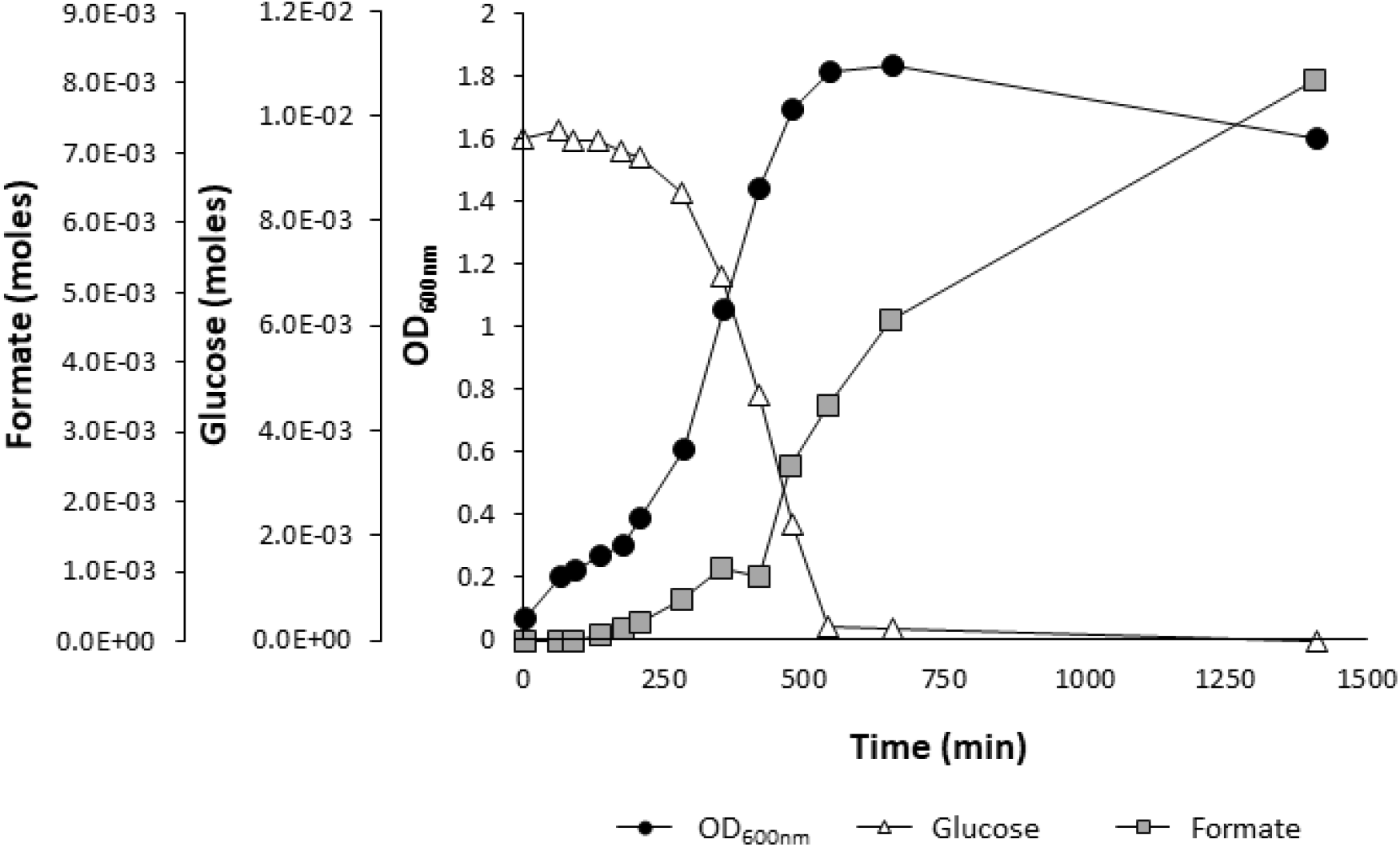
Formate production and bacterial growth profiles of *E. coli* MR60 strain in a pressurised batch reactor. A bioXplorer P400 bioreactor was loaded with rich medium containing 0.8 % (w/v) glucose containing 1 μmol.L^−1^ tungstate salts and operated at 37 °C with a H_2_ and CO_2_ gas sparging flow rate of 50 mL.min^−1^ at ambient pressure. The engineered MR60 strain was inoculated at time point 0 and glucose consumption and formate production was monitored by HPLC.

These results were considered promising since, even at ambient pressure, the performances of the bioprocess using the optimised *E. coli* MR60 strain were considered to outcompete comparable systems in which microorganisms that naturally produce formate, such as *Desulfovibrio sp.,* were used (Mourato et al., 2017). Indeed, while formate produced by the MR60 optimised strain of *E. coli* is in the same range than that produced by *D. desulfuricans*, the maximum rate of formate production is 4-fold higher in the *E. coli* system. Furthermore, while formate production in *D. desulfuricans* started after cell growth cease, formate production started straight upon H_2_ and CO_2_ sparging in the cell culture since the FHL-1 is constitutively express in the optimised strain. Moreover, as formate production is deregulated in this genetic background, formate production continues even after cells entered stationary phase (Figure 4). This clearly emphasizes the potential of an *E. coli* optimised strain for formate production from the hydrogenation of CO_2_ even at ambient pressure.

#### 3.4.2 Effect of elevated pressure on cell growth and formate production by *E. coli*

To investigate the effect of elevated pressure on cell growth, glucose consumption and formate production, the bioreactor was next pressurised at 2, 4 and 6 bar with H_2_ and CO_2_ at a flow rate of 50 mL.min^−1^. Under these conditions, formate production yield (amount of formate produced *per* unit of cell density) could be increased with gas partial pressure up to 4 bar pressure (Figure 5), however no further enhancement of yield of formate produced was observed above 4 bar pressure (Figure 5). Strikingly, however, above 2 bar pressure, the absolute formate content in the bioreactor was seen to decrease drastically (Figure 6C). This was accompanied by a clear inhibition of cell growth under elevated pressures of H_2_:CO_2_ (Figure 6A), and a concomitant drop in glucose consumption commensurate with a low growth rate (Figure 6B). To determine whether the elevated pressure *per se* or the composition of the gas mixture itself was detrimental to the cells, the MR60 strain was subsequently placed in the pressurised bioreactor under 10 bar pressure of 100 % nitrogen gas (Supp. Fig SI S2). Strikingly, neither cell growth rate nor the final cell density was negatively impacted by elevated N_2_ pressure (Supp. Fig. SI S2). This strongly suggests that inhibition of cell growth under elevated H_2_:CO_2_ pressure is likely linked to increasing concentrations of molecular hydrogen and carbon dioxide themselves rather than the elevated pressure alone.

**Figure 5:**
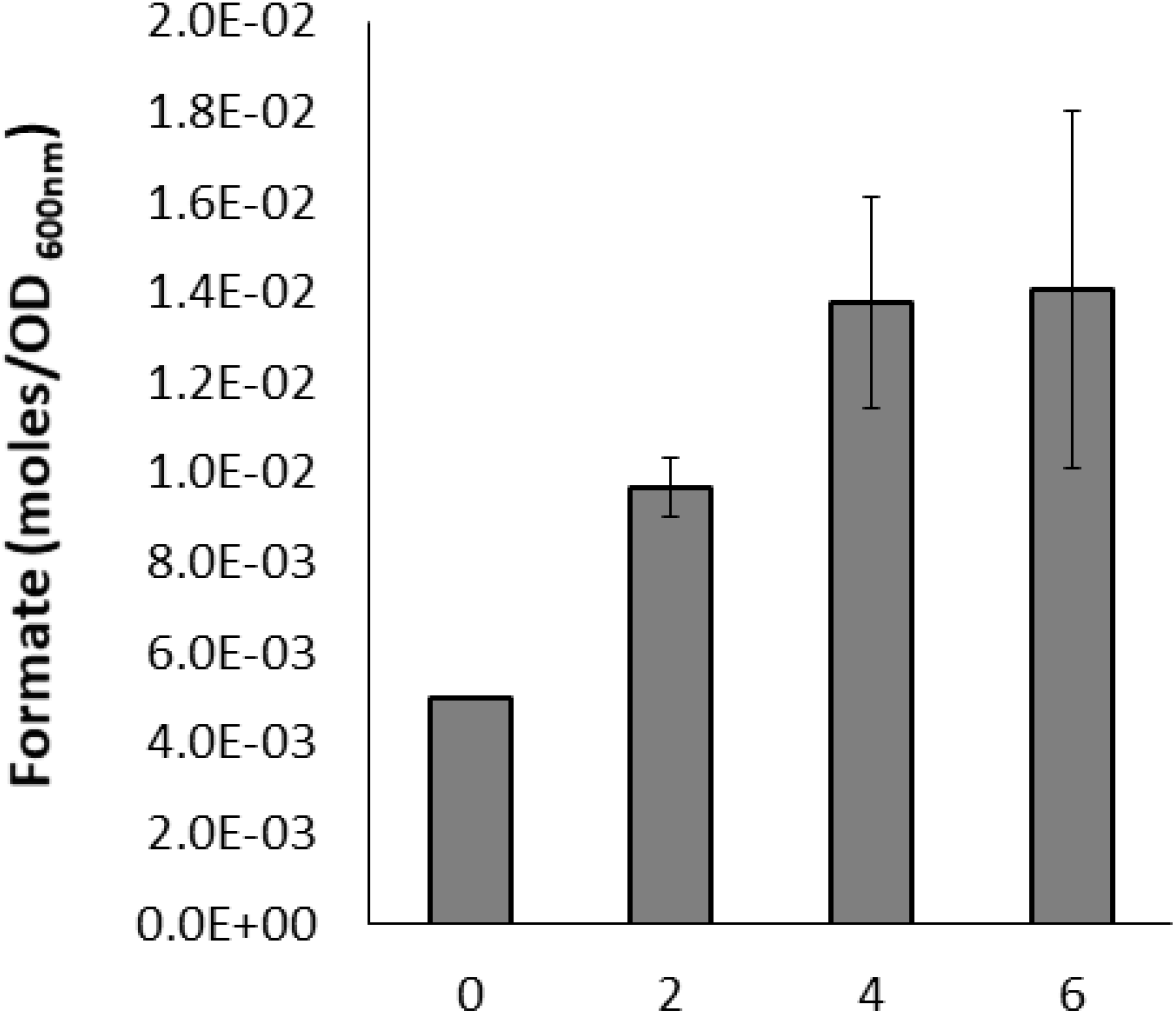
Effect of increasing gas partial pressure on the formate production yield in a pressurized batch bioreactor. A bioXplorer P400 was loaded with rich medium containing 0.8 % (w/v) glucose containing 1 μmol.L^−1^ tungstate salts and operated at 37 °C with a H_2_ and CO_2_ gas sparging flow rate of 50 mL.min^−1^ at ambient (‘0’), 2, 4 and 6 bar pressure. Growth was estimated at the end of stationary phase by measuring the OD at 600 nm. Formate content in the cell suspension was determined by HPLC and yield of formate production calculated considering 1 L culture of *E coli* cells at OD 1 corresponds to 0.25 g_DCW_.

**Figure 6.**
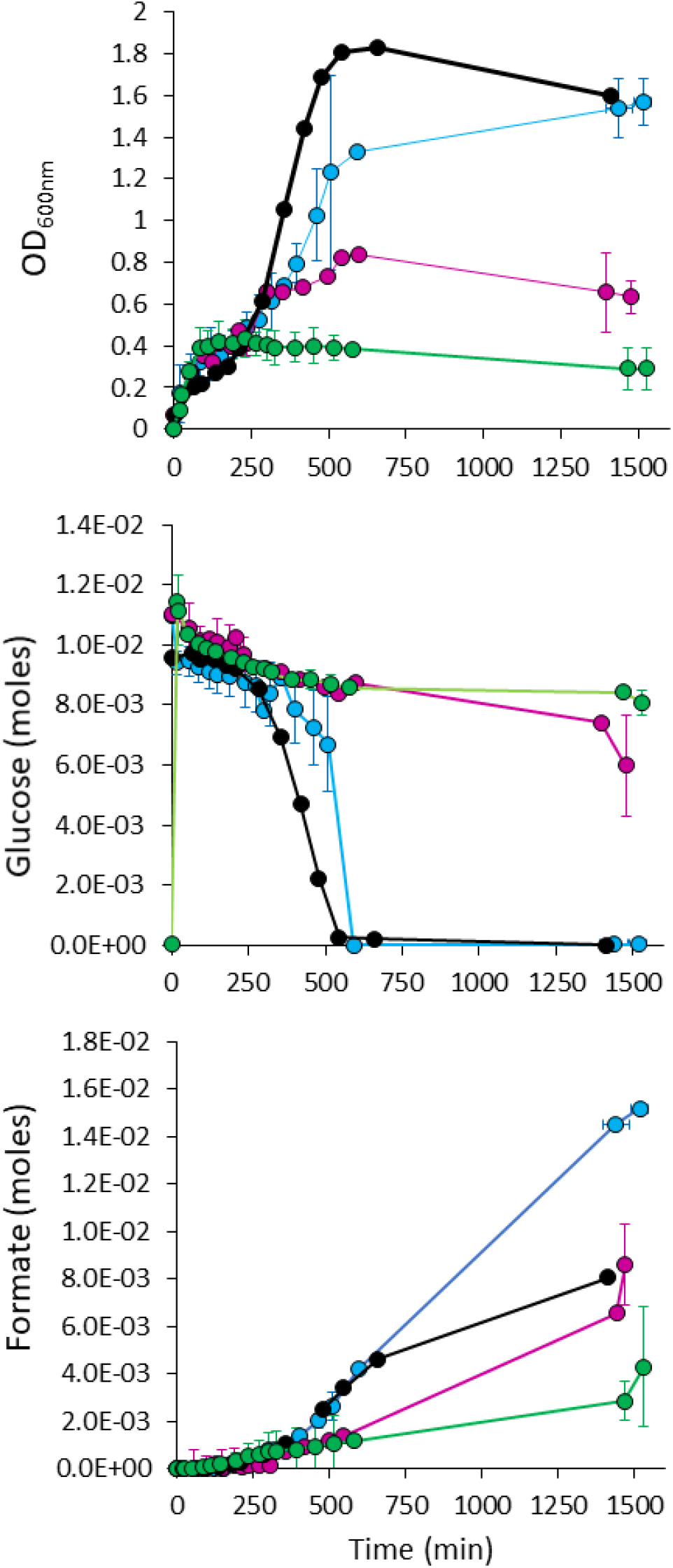
Elevated H_2_:CO_2_ pressure inhibits bacterial growth. A bioXplorer P400 bioreactor was loaded with rich medium containing 0.8 % (w/v) glucose containing 1 μmol.L^−1^ tungstate salts and operated at 37 °C a H_2_ and CO_2_ gas sparging flow rate of 50 mL.min^−1^ at ambient (black), 2 (blue), 4 (purple) and 6 (green) bar pressure. The bioreactor was inoculated with the MR60 strain at time point 0. Growth was monitored at **(A)** OD 600 nm while **(B)** glucose consumption and **(C)** formate content in the culture was determined by HPLC.

It should be considered that the HDCR activity itself is detrimental to cell growth under these conditions. FHL-1 shares a common ancestor with Complex I and as such many thoughts have been put forward on how FHL-1 might contribute to the generation of transmembrane ion gradients or even contrubite to the proton motive force. FHL-1 belongs to the so-called group-4 [NiFe] hydrogenases and is closely related to the complex I (Vignais et al., 2001; Marreiros et al., 2013; Sargent, 2016; Finney and Sargent, 2019). It is often proposed that group-4 hydrogenases couple H_2_ production to the translocation of ions across the membrane - as it was recently demonstrated for the membrane-bound hydrogenase (Mbh) from *Pyrococcus furiosus*(McTernan et al., 2014; Yu et al., 2018). Indeed, the FHL-1 membrane arm exhibits two integral membrane subunits: HycD is predicted to be the membrane anchor for the soluble domain; while HycC is an antiporter-like protein predicted to translocate protons/ions (Schuchmann and Müller, 2013; Ceccaldi et al., 2017; Maia et al., 2017; Müller, 2019; Nielsen et al., 2019). Despite the similarity of the membrane arm of FHL-1 and the Complex I (Schwarz et al., 2018), it is has remained unclear if the E.coli enzyme is capable of translocating ions during disproportionation of formate. In fact, this has been a matter of debate for many years(Bagramyan et al., 2003; McDowall et al., 2014, 2015; Trchounian and Sawers, 2014; Pinske and Sargent, 2016; Petrosyan et al., 2019; Trchounian and Trchounian, 2019) and to-date, no studies convincingly established a contribution to proton motive force generation by FHL-1. The growth defects observed here as the cells are forced to carry out HDCR, however, may be additional circumstantial evidence that the forward reaction contributes to critical energy metabolism.

The pressurised bioreactor experiments also revealed that, in addition to the formate produced from the FHL functioning in reverse mode and lactate which resulted from the central carbon metabolism, detectible traces of acetate was also found in the cell suspension (Supp. Fig. SI S3). Given that acetate is a normally produced under anaerobic conditions from acetyl-CoA, the inactivation of the *pflA* gene product should completely redirect the flux of pyruvate through lactate formation rather than acetate. Strikingly, the yield of acetate also increased with H_2_:CO_2_ gas partial pressure, suggesting that somehow, small amount of CO_2_ might be incorporated through an unknown pathway resulting in acetate production. It was previously proposed that high concentration of fermentation acids limits growth and acetate induces the RpoS regulon associated with entry into stationary phase (Kirkpatrick et al., 2001). Nevertheless, such hypothesis can be excluded since the concentration of formate and acetate in these conditions are far from being toxic.

### 3.5 Whole cell biocatalysis of formate production: What’s next?

Among the organisms that naturally produces formate, acetogenic bacteria are strictly anaerobic bacteria that grow by conversion of syngas (H_2_/CO/CO_2_) to acetate and ethanol through an ancient pathway termed the Wood–Ljungdahl pathway, which combines CO_2_ fixation with the synthesis of ATP (Schuchmann and Müller, 2013; Ceccaldi et al., 2017). Formate is an intermediate in this pathway and in some bacteria such *Acetobacterium woodii*, it can be generated from the direct hydrogenation of CO_2_ catalysed by a soluble enzyme complex termed H_2_-dependent CO_2_ reductase (Schwarz et al., 2018). It consists in a Mo/W-formate dehydrogenase module connected to a [FeFe]-hydrogenase module through two [Fe-S]-containing proteins. In their quest for to find an enzyme with superior catalytic properties, Schwartz and co-workers have isolated a W-containing HDCR from a hyperthermophilic bacterium, *Thermoanaerobacter kuvui* that exhibits remarkable kinetic properties with one of the highest turnover constants for CO_2_ reduction and H_2_ production reported to date (Schwarz et al., 2018; Schwarz and Müller, 2020). Nevertheless, the production of formate using whole cells of *A. woodii* or *T. kuvui* requires to be uncoupled from energy metabolism (to avoid further conversion to acetate). This makes these biocatalysts attractive for enzyme-based applications or intact whole cells catalysis (Schwarz and Müller, 2020). However, a challenging task in the future to fully exploit these organisms as microbial cell factories in fermentation studies, is clearly to improve metabolic capabilities as well as energetic status of the cells (Müller, 2019). Alternatively, sulfate reducing bacteria such as *Desulfovibrio* sp. are metabolically highly versatile since they can use sulfate as terminal electron acceptor in the oxidation of H_2_ or organic compounds, or also produce H_2_ when growing fermentatively in the absence of sulfate and in syntrophy with formate/H_2_-consuming methanogens (Pereira et al., 2008; Plugge et al., 2011; Baffert et al., 2019). They exhibit a hight content and diversity of Mo-formate dehydrogenases, W-formate dehydrogenases (Maia et al., 2017; Nielsen et al., 2019; Niks and Hille, 2019) as well as both [FeFe]- and [NiFe]-hydrogenases (Martins et al., 2016; Baffert et al., 2019) which share the same electron acceptor, the small tetraheme cytochrome *c3*(Matias et al., 2005; da Silva et al., 2012). Hence, it was recently demonstrated that bacterium from the *Desulfovibrio* genus can produce formate from H_2_ and CO_2_ (da Silva et al., 2013; Mourato et al., 2017), involving the periplasmic HydAB [FeFe]-hydrogenase (in H_2_-oxidation mode) and the cytoplasmic Mo-formate dehydrogenase enzyme, both most likely wired *via* the periplasmic tetraheme cytochrome *c3* network (Mourato et al., 2017). Interestingly, it was proposed in this study that *D. desulfuricans* was able to grow during the formate production. As a result, concentration of formate in the bioreactor increased until 64 hours where a maximum steady state value of 30 mM formate production was achieved formate concentration was maintained in the bioreactor over 200 hours. However, although this result was outstanding, the whole bioprocess clearly suffered from low biomass yield (Mourato et al., 2017). By contrast, here we demonstrated that *E. coli* can be harnessed for formate production by turning the native FHL membrane bound enzyme complex in to a HDCR enzyme. The optimised strain itself showed comparable performances than *D. desulfuricans* cells grown in batch reactor. Besides, operating the bioreactor at moderate pressure (*e.g.* 2 bar) H_2_:CO_2_, led to a twice increase in the formate concentration in the cell suspension. This clearly demonstrates the potential of engineering *E. coli* strain as host for bio-based production of formate while fixing CO_2_.

## 4 Conclusions

In this study, an *E. coli* host strain was optimised for the bio-based production of formate from H_2_ and CO_2_ in batch bioreactor. The performance of the bioprocess matches that of enzymes that naturally evolved for CO_2_ reduction. Besides, a bioreactor for batch fermentation process of *E. coli* under pressurised H_2_ and CO_2_ was also designed in this study. Hence, we showed that formate production yield in batch bioreactor, in fact, increases with gas partial pressure. Nevertheless, while *E. coli* cells showed an ability to sustain a pressure up to 10 bar N_2_, at 4 bar pressure of H_2_:CO_2_, cell growth deficiency was observed. Nevertheless, a fair compromise can be made at 2 bar pressure with a twice increased in formate production yield and no detriment on the biomass yield. Hence, capitalizing on the rational design of the host strain combined with an innovative reactor technology, here the potential as *E. coli* host for the bio-based production of formate was demonstrated. This process could serve in the future as a basis for the development of continuous fermentation bioreactor. Furthermore, new biocatalysts with remarkable kinetics properties for CO_2_ reduction, are constantly discovered such as the HDCR from *T. kuvii* (Schwarz et al., 2018), or recently the Mo-Fdh enzyme from *Rhodobacter aestuarii* (Min et al., 2020). Nevertheless, genetic tools in these these organisms are often poorly developed, thus postponing potential applications. By contrast, whole-cell biocatalysis using engineered *E. coli* seems to be more promising method and offers the potential for large-scale and low-cost production (Lin and Tao, 2017). Consequently, building on the host strain optimisation strategy, the incorporation of these candidates in *E. coli* is attractive.

## ACKNOWLEDGEMENTS

This research was funded in the UK in part by Biotechnology & Biological Sciences Research Council (BBSRC) responsive mode award BB/S000666/1, in part by a BBSRC Training Grant (BB/T508743/1) administered by the Industrial Biotechnology Innovation Centre in Scotland (IBIoIC), and in part by an Engineering & Physical Sciences Research Council (EPSRC) CO2Chem Network seedcorn award (all to FS). We are grateful to Dr Alex J. Finney (Newcastle University) and to Drs John H. Allan and Ciarán L. Kelly (Northumbria University) for helpful discussions and advice.

## AUTHOR CONTRIBUTIONS

MR was a Postdoctoral Research Assistant who designed all experiments, analysed data, prepared figures for publication, and wrote the paper. TCR was a postgraduate research student who performed experiments and analysed data. FS conceived the project, assembled the research team, designed the research, supervised the research, analysed data, and wrote the paper.

