## Supplemental Figures for "Harnessing *Escherichia coli* for bio-based production of formate under pressurized H_2_ and CO_2_ gases"

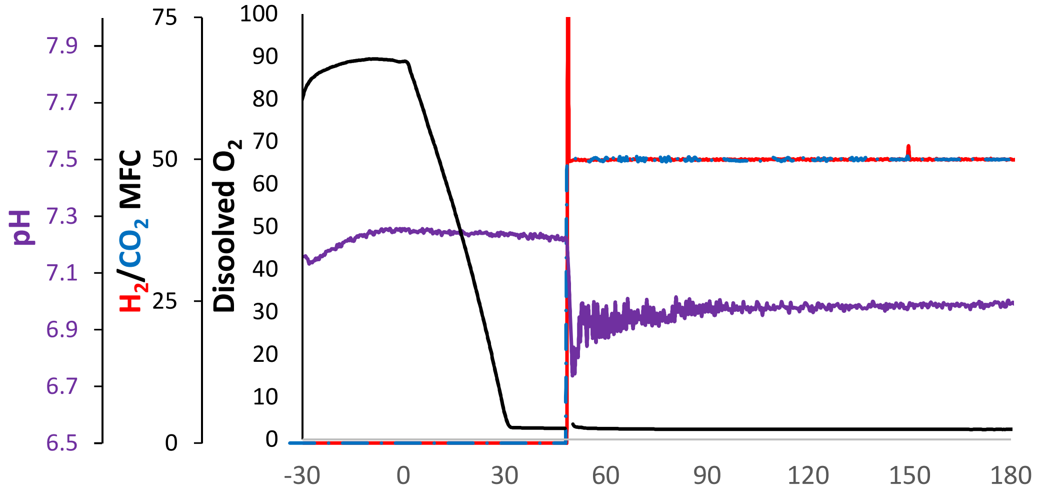


**Supp Figure SI S1:** Batch-fermentation of *E. coli* MR60 strain in the bioreactor at 37°C, 500 rpm, with gas sparging flow rate of 50 mL.min$^{-1}$. The bioreactor was initially fed with rich medium containing 0.8 % glucose supplemented with 1 mmol.L$^{-1}$ sodium tungstate. Cells from an OVN culture were inoculated (T=0 min) at a OD 600 nm of 0.05. Following full consumption of the oxygen initially present in the bioreactor,(black curve), biocatalysis was initiated by sparging $H_2$ and $CO_2$ in the cell suspension (red and blue curves, respectively). This was concomitant with a drop of pH (purple curve) in the cell suspension. As a result, sodium hydroxide was fed to the bioreactor and maintain the pH of the cell suspension at 7 over the time course of the reaction.

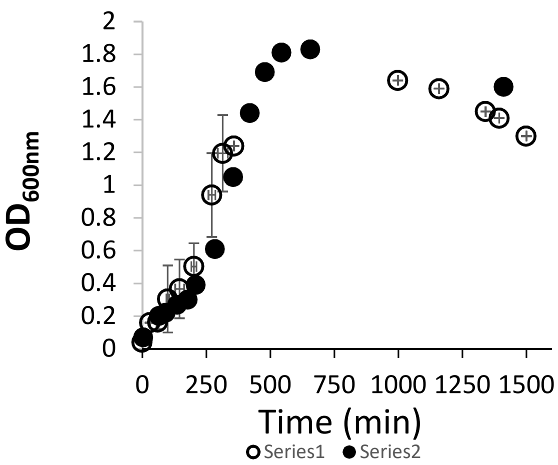


**Supp Figure SI S2:** Effect of elevated pressure on bacterial growth. *E. coli* MR60 strains were grown in the bioreactor at 37 °C with sparging $H_2$:$CO_2$ gas mixture at ambient pressure (in black) or with 100 % nitrogen gas at 10 bars pressure. Growth was monitored at $OD_{600nm}$.

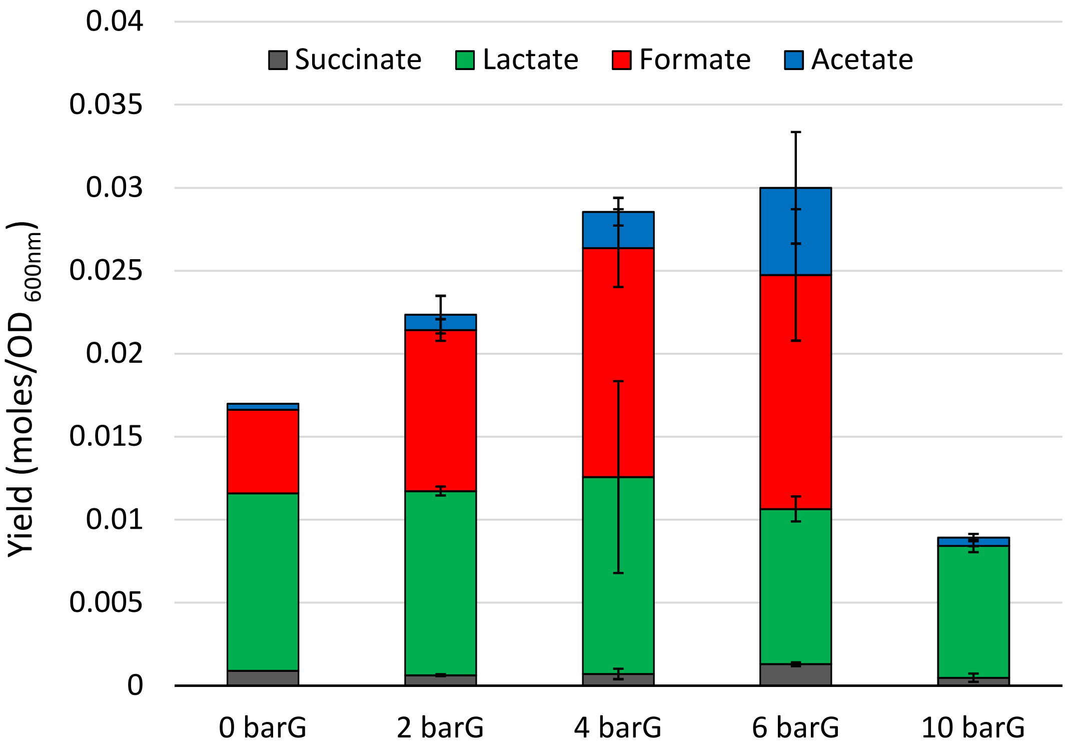


**Supp. Figure SI S3:** Effect of elevated pressure on the metabolism. End-product metabolites produced and secreted by *E. coli* MR60 cells were analysed and quantified by HPLC.
